# MIGS: Methylation Interpolated Gene Signatures Determine Associations Between Differential Methylation and Gene Expression

**DOI:** 10.1101/063941

**Authors:** Christopher E Schlosberg, Nathan D VanderKraats, John R Edwards

**Affiliations:** Center for Pharmacogenomics, Department of Medicine, Washington University in St. Louis School of Medicine, St. Louis, MO, USA

## Abstract

A large number of genomic studies are underway to determine which genes are abnormally regulated by methylation in disease. However, our understanding of how disease-specific methylation changes potentially affect expression is poorly understood. We need better tools to explain specific variation in methylation that potentially affects gene expression in clinical sequencing. We have developed a model, Methylation Interpolated Gene Signatures (MIGS), that captures the complexity of DNA methylation changes around a gene promoter. Using data from the Roadmap Epigenomics Project, we show that MIGS significantly outperforms current methods to use methylation data to predict differential expression. We find that methylation changes at the TSS and downstream ~2kb are most predictive of expression change. MIGS will be an invaluable tool to analyze genome-wide methylation data as MIGS produces a longer and more accurate list of genes with methylation-associated expression changes.

## Introduction

Establishment of specific patterns of DNA methylation is necessary for normal development (1,2), and aberrant methylation is frequently observed in cancer (3,4). CpG islands, regions of high CpG density, are typically unmethylated and associated with 40–70% of mammalian gene promoters (5). Hypermethylation of CpG islands overlapping the transcription start site (TSS) downregulates tumor suppressor genes, thus promoting tumorigenesis (6,7). Typically, promoters are labeled as either methylated and silenced or unmethylated and potentially active (8,9). Although most analysis techniques rely upon this simple binary characterization (10), studies that model methylation using a single window (SW) of ~2kb around the promoter region find only modest negative correlations with expression levels (5,11). Recent work, however, has indicated that a large number of patterns that associate with differential gene expression (12). For example, methylation at CpG island-shores, regions of decreased CpG density flanking CpG islands, correlate with differential gene expression in colon cancer (13). Further, long hypomethylated domains in cancer often contain down regulated genes (13). Positive correlations between gene body methylation and gene expression have also been frequently observed (14,15).

The most common current approach to associate DNA methylation and expression changes is to first identify differentially methylated regions (DMRs) and then associate them with nearby genes. Numerous statistical tools have been developed to identify DMRs (reviewed in (10)). Generally, DMRs are found by segmenting the genome into equally spaced regions and the statistical significance of each region is calculated using a generalized Fisher’s method. To infer biological insight, known genomic regulatory elements are associated with DMRs within a certain distance. However, DMR methods rely on a set of arbitrarily defined thresholds for the size and number of CpGs to include in the DMR. It is often recommended that these parameters be adjusted for each individual dataset, and choices in these parameters can have substantial implications in terms of the numbers of DMRs and genes nearby. However, studies find only weak correlations between DMRs near gene promoters and differential gene expression. One possible reason that both the SW and DMR methods fail to find a strong association between differential methylation and expression is that they attempt to reduce DNA methylation to a single differential value to associate with gene expression change. Neither method considers the surrounding context of these changes.

Here we seek to develop a new approach to model the relationship between changes in DNA methylation and changes in gene expression that accounts for the local methylation context. Our lab previously conducted an unbiased survey of patterns of differential DNA methylation that are putatively associated with gene expression in cellular senescence and upon 5-Azacitidine treatment in Acute Myeloid Leukemia (12, 16, 17). Here we extend this method to classify patterns of methylation change with differential expression. To infer the expression change of novel patterns of differential methylation that may be functionally relevant it would be beneficial to have an algorithm with few assumptions about genic positions, robust in identified regions, and extensible across samples. Therefore, we have developed a supervised classifier, MIGS (Methylation Interpolated Gene Signatures), to identify DNA methylation patterns that associate with transcriptional change. Such a tool will help us understand the functional importance between differential methylation and gene expression.

## Materials & Methods Genome-wide DNA methylation data

Samples from 17 tissues with matched whole genome bisulfite sequencing (WGBS) and RNA-seq were obtained from the Roadmap Epigenomics Project (18). We obtained consolidated methylation data, which was previously cross-assay standardized and uniformly processed. All CpG sites were filtered for ≥4x coverage.

### RNA-seq data

We used uniformly processed protein-coding gene level annotations from Genecode V10 to obtain standardized RPKM values. Each Genecode V10 annotation was converted to Refseq annotations (refGene.hg19.21_12_15.txt) using the myGene.info python API (19). Only genes with unique transcription start sites (TSSs) with complete annotations were considered. All genes with less than 4 exons were removed from analysis for direct comparison with the ROI classifier. If a gene had less than 0.2 methylation change, it was excluded from analysis. All genes shorter than 5 kb (based on genomic distance) were excluded from analysis. If the differentially expressed gene had less than 40 CpGs, it was excluded from the analysis. Differentially expressed genes were defined as genes with ≥2-fold difference between samples after an applied floor of 5 RPKM.

### SW (Single Window)

We compute the average methylation across a single, fixed +/−1 kb window around the TSS of each gene (13). We perform logistic regression using the average methylation in the SW. Logistic regression cross-validation was run with 1000 maximum iterations for the optimization algorithm.

### DMRs (Differentially Methylated Regions)

We use DSS-single to compare DMRs between individual samples (20). We identified DMRs (p<0.01) and used their size (bp), average differential methylation, and stranded distance (bp) to the closest TSS (0 if overlapping) as features for gene expression change classification with a Random Forest classifier with 1001 estimators.

### ROI (Regions of Interest)

The ROI classifier reduces DNA methylation across the entire gene and surrounding regions to multiple averaged values. The ROI classifier was implemented as described in Lou et al. (21).

### MIGS (Methylated Interpolated Gene Signatures)

We applied z-score normalization to each of the CpG values within a 10kb window surrounding the TSS based upon the distribution of methylation values in a 100kb surrounding anchor window. To compare multiple methylation values between different genes, we create methylation signatures using a piecewise cubic hermite interpolating polynomial (PCHIP) to interpolate a curve of z-score normalized differential DNA methylation with a 10kb window around the TSS for each differentially expressed gene. This approach is appropriate since CpG sites exist in different places relative to the TSS of each gene, and CpG methylation values are highly autocorrelated (12, 22, 23). The interpolated curve is then subjected to Gaussian smoothing with a bandwidth of 50bp. To obtain discrete features, we subsample our interpolated methylation signature at 20bp resolution. We then use these features with a Random Forest classifier with 1001 estimators.

### Evaluation Framework

We applied a strict evaluation framework to test the predictions of each method (Figure 2). From 17 samples from the Roadmap Epigenomics Project, we selected 17 pairwise comparisons. All 17 samples were unbiased in the number of CpGs or their mean CpG coverage of the CpGs in the centered 10kb window. We performed 17-fold samplewise cross validation, while holding out any individual samples from the training set that would appear in the testing pairwise comparison. Then for each testing pairwise comparison, we perform 10-fold cross validation for each of the genes in the testing pairwise comparison. To compare between different samples, performance is reported as the accuracy and rejection rate of the number of genes.

**Figure 1:**
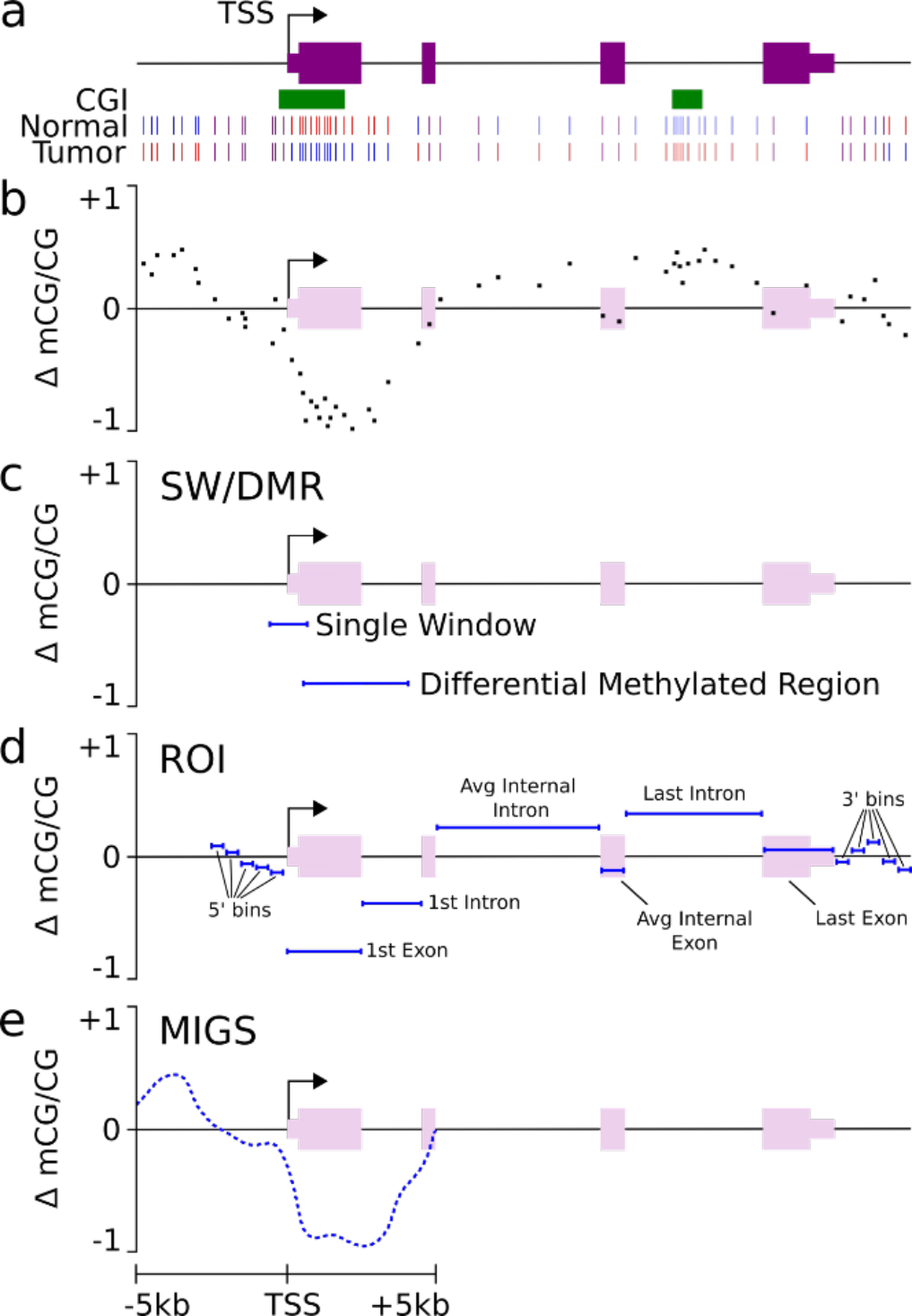
Methods of predicting differential gene expression change from differential DNA methylation. a) Methylation status of an example gene in normal and tumor samples. b) Differential DNA methylation across the example gene. c) Single window (SW) and differentially methylated region (DMR) of the data in (b). d) Region of interest (ROI) representation of the gene in (b). e) Methylation interpolated gene signatures (MIGS) representation of the gene in (b).

**Figure 2:**
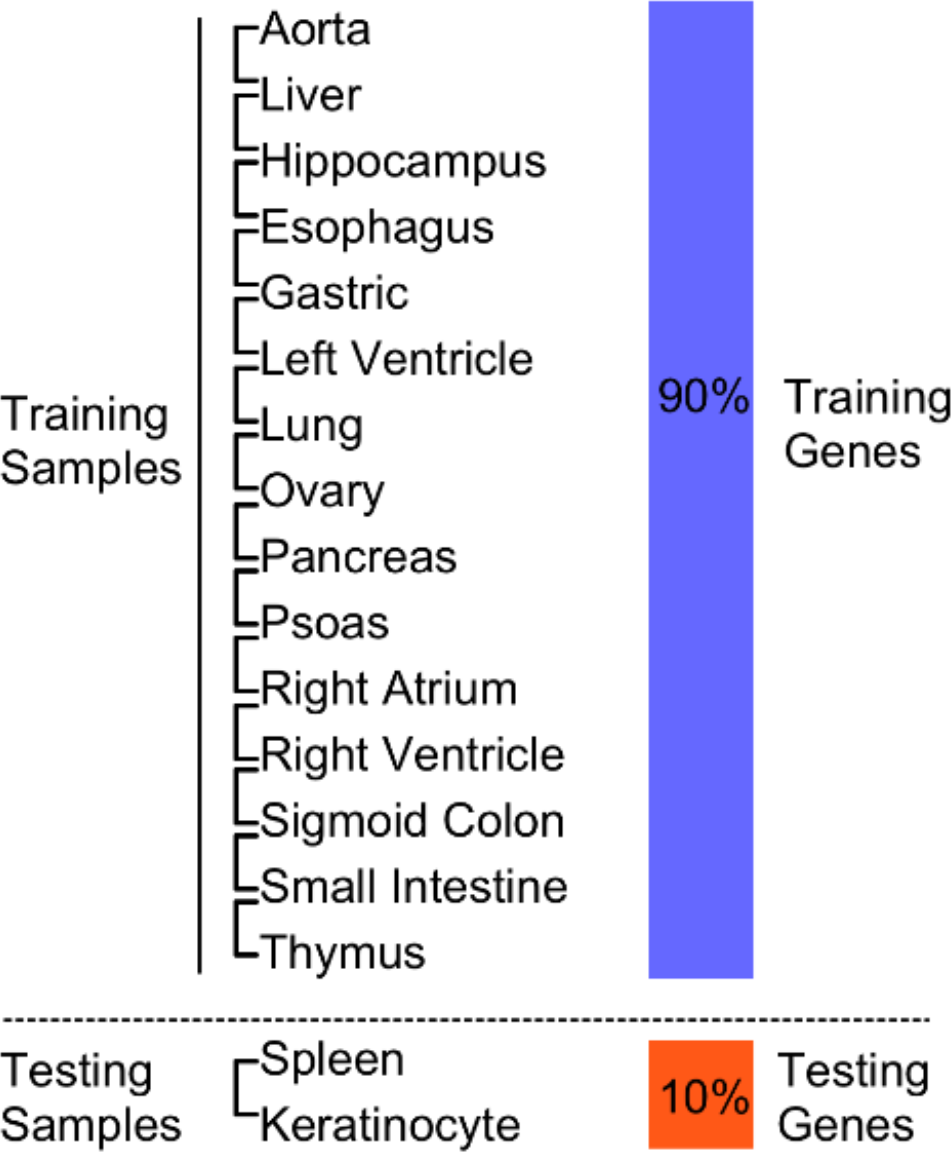
Cross validation comparison framework. Evaluation is performed sample-wise across the 17 samples (left) and then gene-wise across the differentially expressed genes (right).

### Machine Learning Methods

For each of these approaches we applied machine learning methods to classify the direction of differential expression (Table 1). All methods have applied a Random Forest classifier except for the SW method which relies upon logistic regression. Average performance based on the accuracy, reject rate, positive predictive value (PPV), and negative predictive value (NPV) is reported across all genes treated as a large pool from all samples. The ROC curves and the PR curves are averaged to provide a sample-wise level of reporting. All machine learning methods are implemented with scikit-learn and MIGS is publically available on Github at http://github.com/cschlosberg/migs.

**Table 1:** DNA methylation features and classification methods.

| DNA Methylation Representation | Features | Gene Expression Change Classification Method |
| --- | --- | --- |
| Single Window (SW) | $\Delta$ mCG/CG +/-1kb of TSS | Logistic Regression (LR) |
| Differentially Methylated Regions (DMR) | Distance from TSS to DMR (bp)<br>DMR width (bp)<br>Avg. $\Delta$ mCG/CG | Random Forest (RF) |
| Regions of Interest (ROI) | Avg. $\Delta$ mCG/CG:<br>Five 400bp bins 5' of TSS<br>1<sup>st</sup> Exon<br>1<sup>st</sup> Intron<br>Avg. Internal Exon<br>Avg. Internal Intron<br>Last Exon<br>Last Intron<br>Five 400bp bins 3' of txEnd | Random Forest (RF) |
| Methylation Interpolated Gene Signatures (MIGS) | $\Delta$ mCG/CG of 500 bins (20bp) +/-5kb of TSS | Random Forest (RF) |

## Results

### MIGS

MIGS is built as a supervised framework using a spatial representation of DNA methylation surrounding the TSS. We model methylation using a signature in the +/− 5kb window around the TSS. This signature allows the model to incorporate the entire profile of methylation changes across the area in and around the gene’s promoter including any associate CGI and CGI-shore regions. Interpolation and smoothing of the data decreases the influence of sequencing error at individual CpGs (12). Similar approaches have shown a marked improvement in the ability to determine DMRs (24). Since the goal of most studies is to identify how changes in methylation alter expression in a disease, we examine the ability of methylation to predict differential expression change. The Random Forest classifier provides a nonparametric model of expression change classification with a low number of hyperparameters, generation of an internal unbiased estimate of testing error, and identification of feature importance (25).

### Implementation of Other Methods

To understand the performance of MIGS we compared it to the most common methods used in the literature (Figure 1). The first method is to model methylation using a single window (SW) +/− 1 kb window around the TSS, and use logistic regression to predict differential expression based on the average methylation in this window. The second method is a DMR approach. Each gene was first associated with its closest DMR as identified by DSS (20). We extracted features including the distance from the DMR to the gene, the average methylation over the DMR, then associate the DMR with the closest gene. These features were then used in a Random Forest classifier to predict differential expression. The third method was to implement the ROI (Region of Interest) classifier for differential expression. The ROI, originally developed for use on individual samples, uses the average methylation across bins at and nearby the gene’s promoter and exon and intron boundaries as features for a Random Forest to predict expression (21).

### MIGS best predicts gene expression change from differential methylation

To compare classifiers we used whole genome bisulfite sequencing (WGBS) DNA methylation and RNA-seq data from the Roadmap Epigenomics Project for 17 tissue samples (18). We have implemented a sample-wise and 10-fold gene-wise cross validation framework to ensure that each testing gene does not observe an example of either its own gene or sample (Figure 2). Since patterns of differential DNA methylation can be very similar between datasets, this evaluation framework tests the strength of the DNA methylation representation and the universality of DNA methylation patterns rather than the ability to simply recall an observed gene’s expression class. Since transcription can be controlled by multiple factors other than DNA methylation, we introduce a reject rate. The reject rate excludes genes that cannot be reliably predicted (i.e. they likely do not have methylation-associated expression changes). Each classifier provides a probability of classification, which can be used as a threshold to compute the rejection rate.

In Figure 3a, we observe that MIGS outperforms ROI, SW, and DMR methods in accuracy and proportion of the genes returned (1-Reject Rate) across all rejection rates. Importantly, MIGS returns many more genes more accurately at high levels of its probability of classification. MIGS further outperforms the other methods when classifying the entire dataset by ROC and precision-recall analysis (Figure 3b,c). We next examined whether there was any implicit bias between the expression classes called at each probability of classification for each method. We observe that MIGS does not appear to have any bias towards the positive (up-regulated) or negative (down-regulated). In contrast, there appears to be a slight bias in favor of the positive class for the ROI, SW, and DMR methods at higher probabilities of classification (Figure 3d,e).

**Figure 3:**
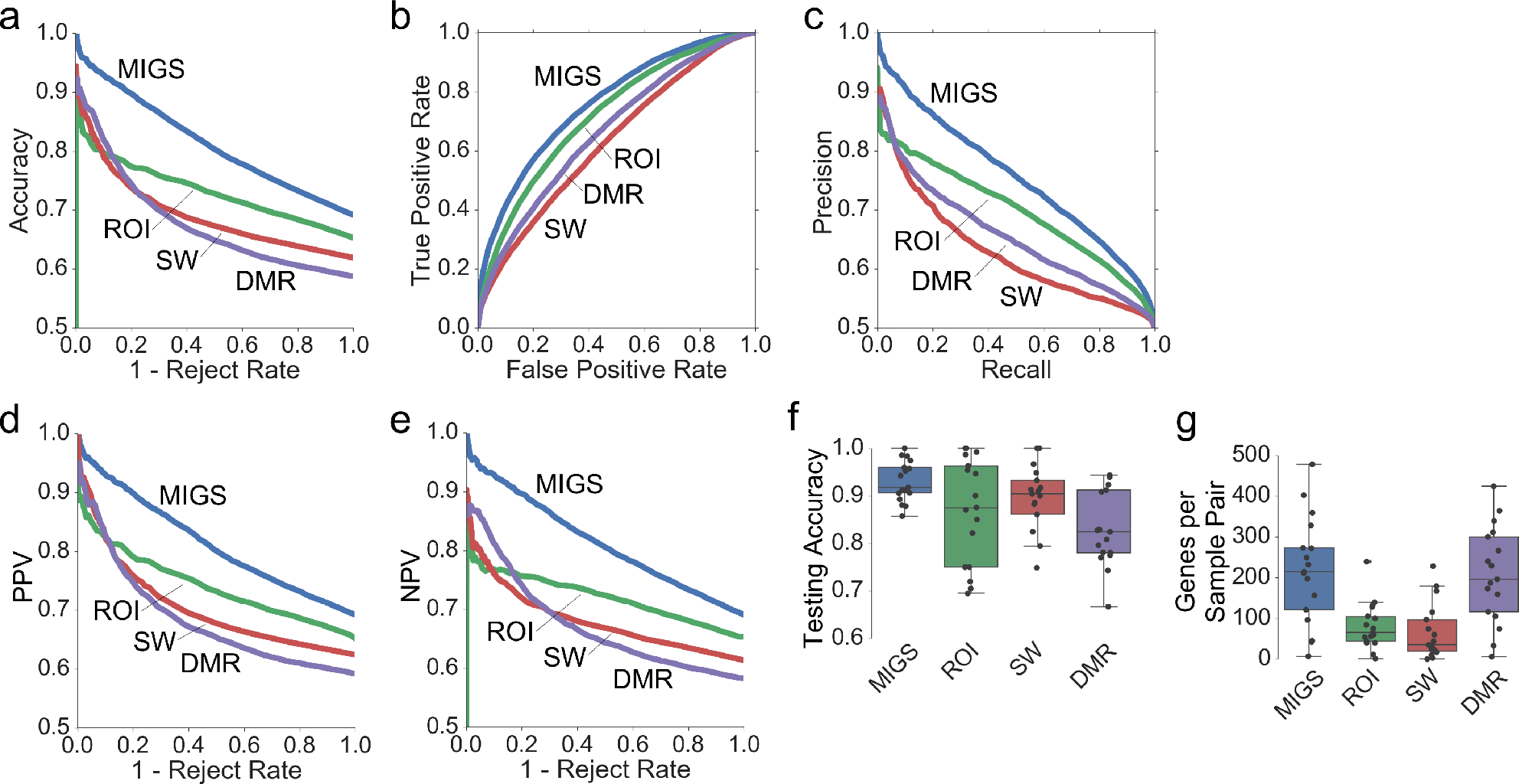
Evaluation of each classifier using 17 tissue samples. a) Accuracy versus 1-Reject Rate b) ROC curve c) PR curve d) PPV versus 1-Reject Rate e) NPV versus 1-Reject Rate f) testing accuracy at 90% operating probability of classification g) number of genes returned at 90% operating probability of classification.

While it is important to examine the performance of each classifier over a range of probabilities of classification, in practice, we would set an operating parameter to examine the accuracy and the number of genes returned. Since it is unclear what threshold to set for the probability of classification when examining a new dataset, we examined the performance of accuracy versus probability of classification for each sample. MIGS matches the accuracy given the probability of classification indicating that this probability can be used as an estimate of the final classification error in cross-sample comparisons. At an operating probability of classification at 90%, we observe that MIGS returns the largest number of genes (Figure 3g), and that those genes are returned at the highest level of accuracy (Figure 3f). Each of these evaluation metrics in our framework indicate that MIGS provides the most optimal representation of DNA methylation in comparison to the methods analyzed.

### 3’ proximal and TSS regions are most predictive of differential expression

We next examined which features are most important for classification. We first developed a series of MIGS classifiers each using signatures from 5kb windows centered at varying distances away from the TSS (Figure 4a,b). Performance peaks for a window centered 2kb downstream of the TSS, indicating that the most important features exist downstream of the TSS. Performance only moderately improved for the entire 10 kb window compared for the best 5kb window. An analysis of feature importance in the Random Forest using a classifier trained from all 17 samples showed that the most important features for gene expression classification occurs downstream ~1–2kb of the TSS with a minor contribution centered on the TSS and ~0.5kb upstream (Figure 4e).

**Figure 4:**
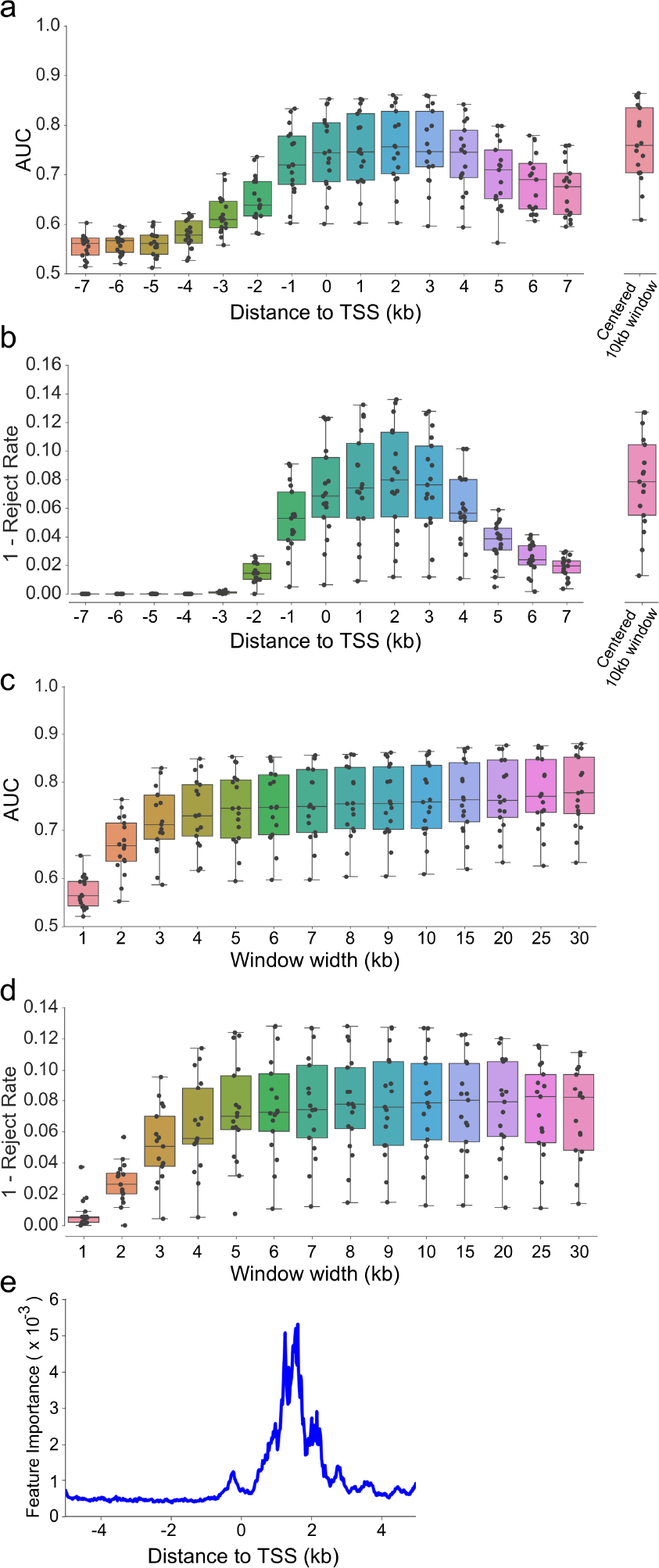
DNA Methylation downstream of TSS is important for classification. a) ROC curve AUC and b) 1-Rejection Rate for MIGS methylation signatures created for 5 kb windows centered at varying distances to TSS. c) ROC curve AUC and d) 1-Rejection Rate for MIGS methylation signatures created using varying window sizes each centered at the TSS. e) MIGS RF feature importance.

We next designed a series of MIGS classifiers to examine how the size of the centered window affects performance, with constant 20bp subsampled resolution. Increasing the window size showed steady improvements in performance. However, after the window size increases greater than 10 kb there is a substantial loss in the number of genes returned, which can be attributed to increased noise from DNA methylation in regions distal to the TSS (Figure 4d).

## Discussion

MIGS outputs a ranked gene list to identify genes where there exists a putative association between methylation and expression. At any probability of classification, MIGS returns the largest numbers of genes at the highest level of accuracy. The SW can achieve high levels of accuracy, but only a much lower number of genes. Conversely, the DMR method generally returns a low level of accuracy with a much higher level of genes. DMRs require a large amount of tuning to find appropriate genome segmentation parameters, which implies that the nearest DMR might not necessarily be the most informative DMR to predict the expression class. The ROI classifier returns a relatively low number of genes with a larger variation in the accuracy of the genes returned.

When applied to expression in a single sample, bins around the TSS were most important for classification. Incorporating the complexity of patterns rather than reducing methylation to a single or even multiple averaged values is critical for the success of MIGS. Further, interpolation and smoothing serve to decrease inaccuracy of low coverage methylation calls, as has been observed in DMR callers. MIGS performance demonstrates that full methylation profiles across the TSS region are most predictive of differential gene expression. The association of methylation and expression downstream of the TSS agrees with multiple other cancer-and tissue-specific studies (26–30). Given differential methylation data, MIGS takes full advantage high resolution DNA methylation data to provide accurate predictions and probabilities of expression change.

## Acknowledgements

We would like to acknowledge Tao Ju for providing assistance on best practices for feature representation and Kilian Weinberger for confirmation of evaluation framework of our machine learning methods.

